# Exploring the Structural Lexicon of the Proteome via Metric Geometry

**DOI:** 10.1101/2025.11.03.686256

**Authors:** Elijah Gunther, Pablo G. Camara

## Abstract

The three-dimensional structure of proteins is intimately linked to their function, yet establishing comprehensive frameworks for systematically comparing and organizing protein structures across the proteome remains a significant challenge. Here, we introduce GWProt, a computational framework that leverages recent advances in metric geometry, such as Gromov-Wasserstein couplings, for protein structure alignment and analysis. GWProt enables the integration of biochemical information into structural comparisons and introduces the concept of local geometric distortion, a measure that captures local conformational differences. We demonstrate the utility of this framework by identifying conformational switches within individual proteins, detecting functional domains shared among evolutionarily distant viral proteins, revealing topological rearrangements in homologous folds, and uncovering recurrent short structural motifs underlying functional domains across the human proteome. Collectively, these results establish the use of metric geometry as a versatile and quantitative framework for the systematic comparative analysis of protein structures, complementing existing approaches for elucidating protein organization.

## Introduction

Proteins are the fundamental information-processing units of all living systems, with their functional capabilities determined by the dynamic 3D conformations encoded in their amino acid sequences. Understanding the diversity and governing principles of protein structure is therefore of central importance, and substantial efforts have been devoted over the past four decades to developing algorithms for protein structure alignment^1^, yielding widely adopted methods such as DALI^2^, SSAP^3^, CE^4^, and, more recently, TM-align^5^, Foldseek^6^, and Reseek^7^. The relevance of these approaches has grown in parallel with the rapid expansion of the Protein Data Bank^8^ (PDB), which now houses hundreds of thousands of experimentally determined protein structures, and has been further amplified by recent advances in deep learning that enable accurate prediction of protein structures directly from sequence^9–11^. These developments have highlighted the modular design of proteins and facilitated the identification and systematic classification of functional protein domains^12–16^. Additionally, they have led to the recognition of smaller recurrent building blocks of protein architecture, such as super-secondary structural elements^17–21^ and 3D microenvironments^22–24^, although their systematic classification remains incomplete, partially due to the lack of a unifying organizational framework.

From a mathematical standpoint, the study and comparison of geometric structures fall within the discipline of metric geometry^25^. The primary objects of study in metric geometry are metric spaces: collections of points equipped with a notion of distance between them. Since the introduction of the modern concept of metric space by Maurice Fréchet in the early twentieth century^26^, the field has undergone several major developments. In 1981, Mikhail Gromov transformed it with the introduction of the Gromov-Hausdorff distance^27^, which provided a rigorous framework for quantitatively comparing metric spaces independently of rigid transformations such as rotations and translations. However, because computing the Gromov-Hausdorff distance involves solving an optimization problem of exponential complexity in the number of points, its impact initially remained largely theoretical, with few practical applications. Over the past decade, this has changed with the development of rigorous and computationally efficient relaxations, such as the Gromov-Wasserstein distance^28–30^, which reformulates the problem within the framework of optimal transport theory and can be approximated in nearly linear time^31–33^. These advances have established a mathematical foundation for the quantitative study and comparison of geometric structures encountered in real-world applications, including cell morphometry^34^, single-cell and spatial omics^35–39^, neuroscience^40^, and communication networks^41^.

In this work, we investigate the application of these advances in metric geometry to the problem of protein structure alignment and extend them by introducing the notion of local geometric distortion. Our results demonstrate that metric geometry provides a powerful and versatile framework for the systematic comparative analysis of protein structures, complementing existing approaches with several strengths, such as the ability to detect conserved structural motifs, the capacity to incorporate biochemical information into structural comparisons, and robustness to topologically non-trivial sequence rearrangements. We demonstrate this framework with the analysis of evolutionarily distant viral proteins, and apply it to the human proteome, where we show that metric geometry can serve as an organizing principle for the systematic study of recurrent structural polypeptide fragments.

To facilitate use, we have implemented and documented the computational methods of this work as an open-source software package, available to the broader community. Together, our results establish the use of metric geometry as a complementary quantitative framework for advancing the understanding of protein organization, function, and evolution.

## Results

### GWProt: Gromov-Wasserstein correspondences for protein structural alignment

The problem of establishing a local correspondence between two shapes, or *metric spaces*, is a central problem in metric geometry. The Gromov-Hausdorff distance between two metric spaces establishes an optimal correspondence between their points by minimizing a loss function that quantifies the total geometric distortion induced by the mapping^28,30^ (Fig. 1A). The value of the loss function at its minimum defines a structural distance, satisfying the triangle inequality and all other mathematical properties required of a distance function. Intuitively, the Gromov-Hausdorff distance captures the minimal deformation needed to transform one shape into another. Thus, it provides a natural and rigorous mathematical framework for structurally aligning proteins, allowing us to determine optimal residue-to-residue pairings between two protein structures (Fig. 1B).

**Figure 1.**
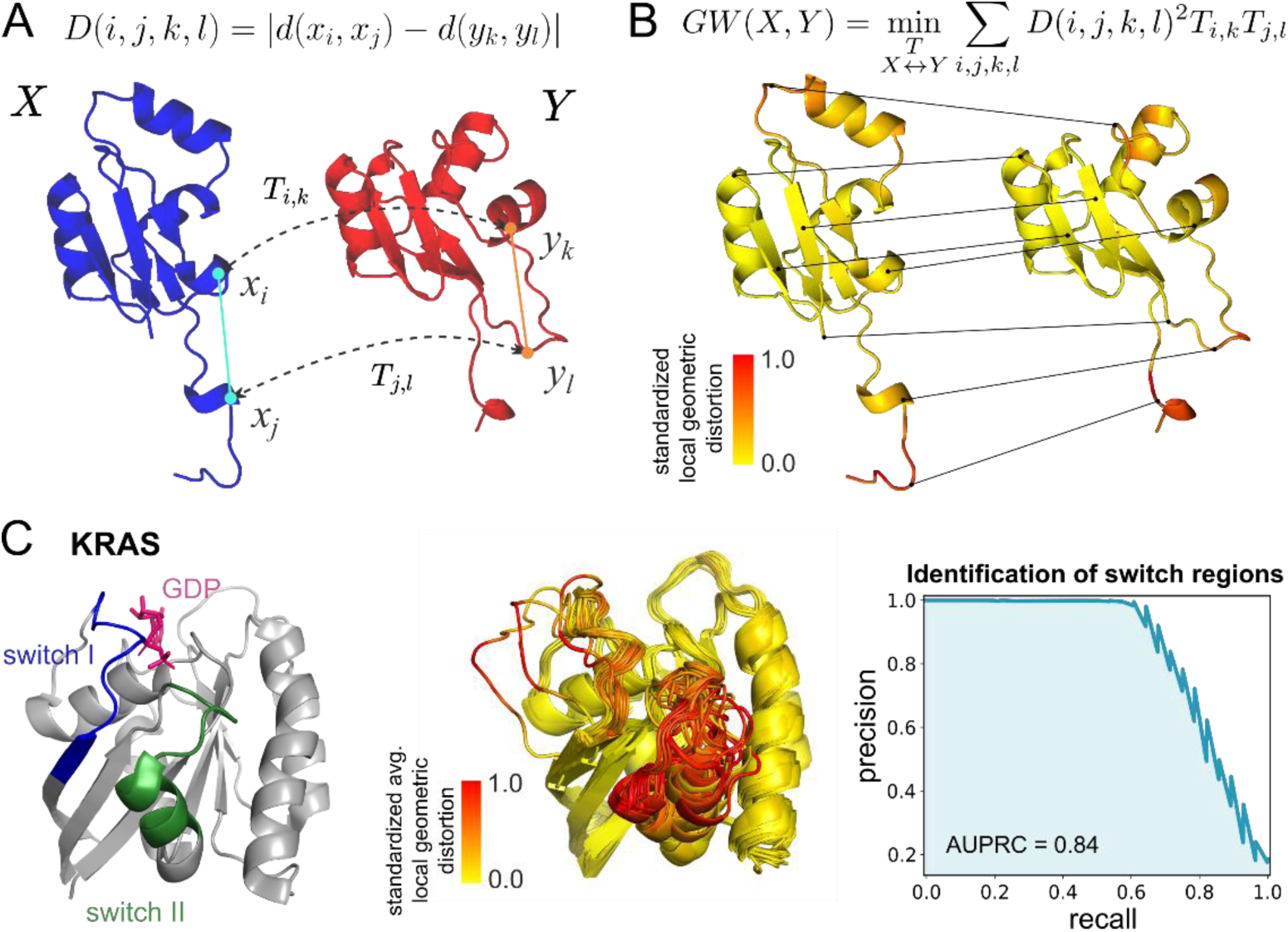
Gromov-Wasserstein correspondences for protein structural alignment. **A)** Schematic illustrating the concept of geometric distortion underlying Gromov-Hausdorff and GW distances. Given a correspondence 𝑇 between two protein structures 𝑋 and 𝑌, the geometric distortion 𝐷(𝑖, 𝑗, 𝑘, 𝑙) between a pair of C_α_ atoms 𝑥_𝑖_ and 𝑥_𝑗_ in protein 𝑋 and their matched α-carbons 𝑦_𝑘_ and 𝑦_𝑙_ in protein 𝑌 is defined as the discrepancy between the Euclidean distances 𝑑(𝑥_𝑖_, 𝑥_𝑗_) and 𝑑(𝑦_𝑘_, 𝑦_𝑙_). **B)** The GW correspondence between two protein structures is the mapping 𝑇 that minimizes the total geometric distortion. The local geometric distortion quantifies the contribution of each C_α_ atom to the optimal cost 𝐺𝑊(𝑋, 𝑌). In the figure, the optimal GW correspondence is visualized for two proteins, with colors indicating local geometric distortion, showing regions of high (red) and low (yellow) structural agreement. **C)** Identification of switch regions in KRAS. Left: Backbone of the KRAS protein with annotated switch I and switch II regions. Middle: GW-based alignment of 54 experimentally determined KRAS structures, colored by average local geometric distortion. The switch regions show high structural variability. Right: Median precision-recall curve for a predictor of switch regions based on the average local geometric distortion. AUPRC: area under precision-recall curve.

The computation of Gromov-Hausdorff correspondences is an NP-hard problem, making it computationally unfeasible for most practical applications. However, efficient relaxations can be achieved by using probabilistic correspondences, as they enable reformulating the problem in terms of optimal transport theory^28–30^. The Gromov-Wasserstein (GW) distance approximates the Gromov-Hausdorff distance and provides a measure of structural similarity by identifying an optimal weighted correspondence between the two shapes (Fig. 1B, Methods). Like the Gromov-Hausdorff distance, the GW distance is a true distance function.

The computation of GW correspondences (more commonly known in the mathematical literature as GW couplings) is a non-convex optimization problem, for which fast algorithms and near-linear runtime approximations have been developed^32,42–44^. Furthermore, recent extensions, such as fused and unbalanced variants^40,45,46^, enable the integration of additional features in the correspondences (e.g., local biochemical properties) and support partial structural alignments.

To investigate the utility of GW correspondences for protein structure alignment, we developed a Python package, GWProt, which computes GW and fused GW couplings between the sets of C_α_ atoms of protein pairs. Additionally, by expressing the GW distance as a sum over individual residue contributions, we introduced the concept of local geometric distortion, which quantifies the degree of structural conservation of each residue in the alignment (Fig. 1B, Methods). GWProt is freely available to the community as open-source software (see Code Availability).

### Local geometric distortion identifies active sites in the KRAS oncoprotein

As an initial proof of concept to evaluate the utility of GW correspondences for structural protein alignment, we analyzed 54 crystallographic structures of the human GTPase KRAS (Kristen Rat Sarcoma), including both wild-type and mutant forms, determined by X-ray diffraction and obtained from the RCSB Protein Data Bank^8^ (Supplementary Table 1).

KRAS cycles between an inactive guanosine diphosphate (GDP)-bound and an active guanosine triphosphate (GTP)-bound state. In its GTP-bound form, it binds and activates various effector proteins within the RAS signaling pathway. The interface for effector binding is formed by two flexible regions, known as switch I and switch II^47^ (Fig. 1C).

We used GWProt to compute GW correspondences between all KRAS structures (including both GDP- and GTP-analogue-bound forms), as well as the local geometric distortion at each C_α_ atom (Fig. 1C). As expected, the resulting correspondences were close to the identity map (Pearson’s correlation with the identity map, 𝑟 = 0.985 ± 0.021), reflecting the high degree of homology among the crystallographic structures. Moreover, most of the local geometric distortion was concentrated in the two switch regions. Thus, using average distortion values alone, we were able to predict the location of the switch sites in each structure with an area under the precision-recall curve (AUPRC) of 0.84 ± 0.03 (Fig. 1C).

These results demonstrate that GW correspondences can efficiently capture local structural variation and identify conserved and variable regions across closely related protein conformations.

### Local geometric distortion identifies functional domains in evolutionarily distant viral proteins

Having assessed the utility of GW correspondences for identifying structurally variable regions among closely related protein conformations, we next turned our attention to identifying structurally conserved regions among distantly related proteins.

For that purpose, we analyzed the computationally predicted structures of the core domain of 2,777 RNA-dependent RNA-polymerases (RdRps) from Riboviruses spanning 21 taxonomic classes^48,49^ (Supplementary Table 2). The RdRp core domain binds the RNA strand and comprises palm, fingers, and thumb subdomains^50^, arranged in a fashion analogous to a hand gripping the strand. The palm subdomain contains the catalytic motifs A, B, and C, which are highly conserved among most known RdRps^51^ (Fig. 2A). However, although the palm subdomain is structurally conserved, its amino acid sequence identity can be as low as 10%^52^.

**Figure 2.**
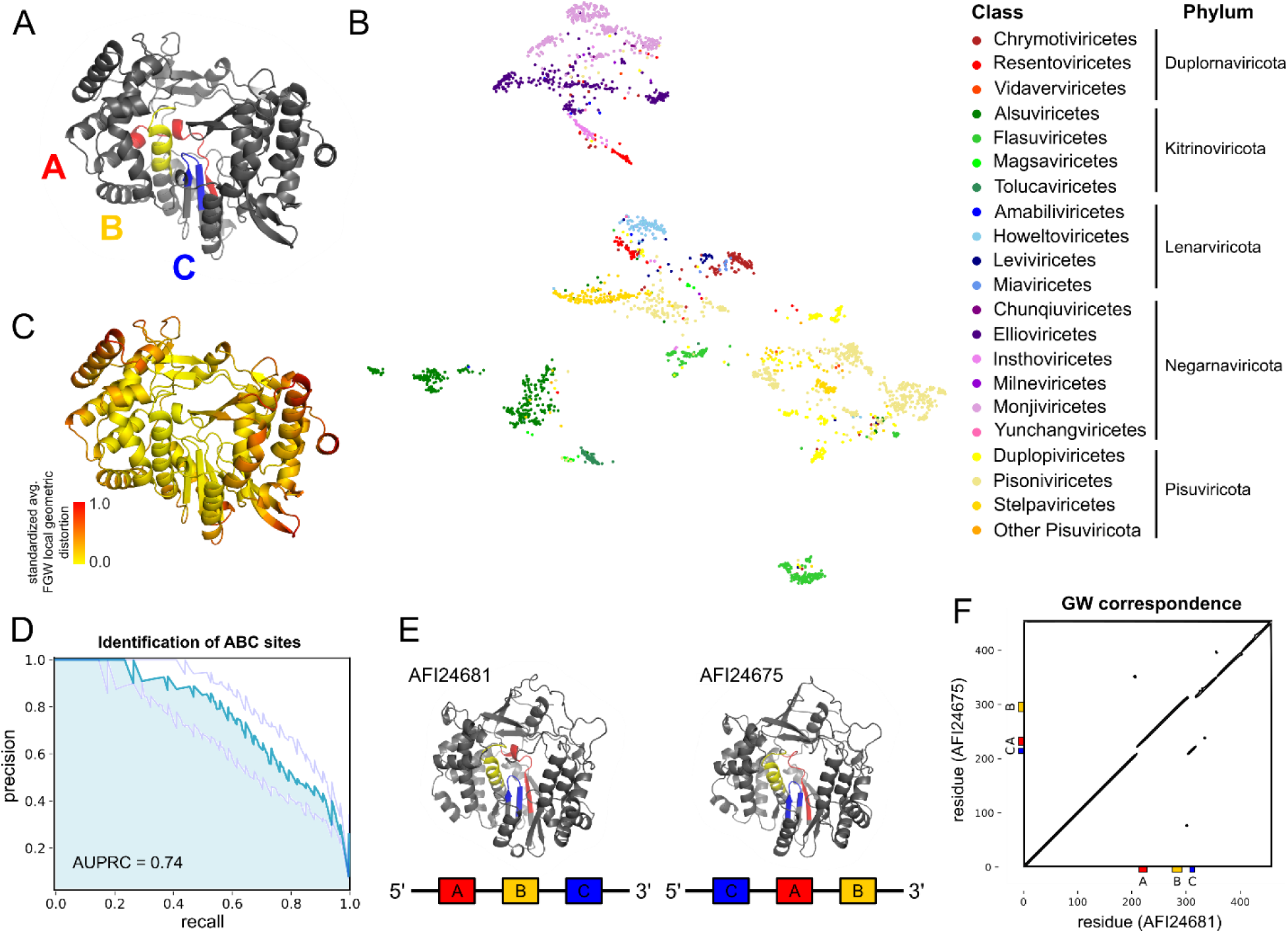
Identification of structurally conserved catalytic sites in RdRps. **A)** Hepacivirus hominis (GenPept ID AFD18577) RdRp core domain with A, B, and C motif sites highlighted. **B)** UMAP representation of the GW-based structural space of 2,777 RdRps. The UMAP is labeled by Ribovirus class, showing consistency between structural organization and viral taxonomy. **C)** Example of an RdRp core domain colored by average FGW local geometric distortion. Regions of low distortion, mostly located at the catalytic core, are structurally conserved. **D)** Median, 20%, and 80% percentile precision-recall curves for predicting A, B, or C sites based on average FGW local geometric distortion across 97 randomly selected RdRps. AUPRC: area under the precision-recall curve. **E)** Example of ABC and CAB RdRp core domains showing a high degree of structural homology. **F)** Dot plot visualization of the GW correspondence between the two proteins shown in (E). The correspondence clearly captures the genomic circular permutation between the A, B, and C motifs.

We aligned the RdRp structures using GW correspondences and visualized the resulting structural distances using Uniform Manifold Approximation and Projection^53^ (UMAP) (Fig. 2B). In this representation, RdRps from different viral classes clustered distinctly, supporting the use of RdRp protein structures for the taxonomic classification of RNA viruses that lack a DNA stage of replication^48,49^. To quantify the consistency of the GW structural space with the established taxonomic classification of Riboviria, we trained a *k* = 3 nearest-neighbor classifier to predict viral class based on the GW distances to RdRp structures in the training set, achieving a Matthew’s correlation coefficient (MCC) of 0.94 (10-fold cross-validation).

We used the average local geometric distortion to identify structurally conserved regions in 97 randomly selected RdRps (Methods). The inner region, corresponding to the catalytic core, exhibited substantially lower distortion than the outer regions (Supplementary Fig. 1), indicating higher structural conservation. For each protein, we then computed a precision-recall curve to assess the accuracy of predicting the A, B, and C motif sites based on regions of low local geometric distortion (Supplementary Fig. 1), using the output of Palmscan^48^, a sequence-based software specifically designed to identify A, B, and C motifs, as the ground truth. The average AUPRC from this analysis was 0.72 ± 0.11, indicating a good degree of agreement between A, B, and C sites and low-distortion regions. However, the accuracy of this analysis is likely constrained by the presence of other structurally conserved regions, such as D and E motifs, which are not detected by Palmscan.

We then reasoned that incorporating biochemical information into the computation of GW correspondences and local geometric distortion could enhance the identification of functional sites. To test this, we recomputed the alignments using a fused GW approach incorporating the hydrophobicity of each residue in the computation of correspondences (Methods). This modification of our approach led to an improvement in the accuracy of A, B, and C site predictions, with an average AUPRC of 0.74 ± 0.11 (Figs. 2C and D; Wilcoxon signed rank test p-value = 3 x 10^-14^).

Collectively, these results demonstrate the utility of local geometric distortion as an unsupervised and interpretable method for identifying structurally conserved regions in evolutionarily distant proteins and show that this signal can be further enhanced by integrating orthogonal biochemical information into the computation of GW correspondences.

### GWProt structural alignment detects internal permutations in homologous proteins

Structural rearrangements facilitate the modular evolution of protein domains, with structural motifs often recurring in different combinations across proteins. Some of these rearrangements do not preserve the sequential order of motifs, with circular permutations of the entire protein being the most prominent example^54^. In certain cases, however, topologically non-trivial rearrangements affect only a small subset of the protein. For instance, in the RdRps of several ribovirus families, the conserved A, B, and C motifs appear in CAB order in the amino acid chain, rather than the canonical ABC sequence^55,56^ (Fig. 2E). Detecting such internal permutations poses a significant challenge, as most structural alignment methods rely on sequential ordering or are tailored to specific cases, such as circular permutations^57^ or permuted RdRps^48^. Only a few general ordering-independent methods based on computer vision approaches like geometric hashing have been developed to address this problem^58–61^.

Since GW correspondences minimize the geometric distortion between pairs of C_α_ atoms, they are independent of the sequence order, and the resulting structural alignments should be robust to internal permutations. To verify this, we examined the GW correspondences between the catalytic cores of ABC and CAB RdRps with high structural homology. As anticipated, the correspondences unequivocally identified the permutation of active sites and were consistent with maps produced by existing ordering-independent methods (Fig. 2F and Supplementary Fig. 2), confirming the robustness of GW correspondences to complex amino acid sequence rearrangements of structurally conserved motifs.

### GW distances enhance the discovery of previously unseen viral phyla

The value of the GW cost function at its minimum defines a distance function that can be used to quantify protein structure similarity^29^. Although numerous algorithms for protein structural similarity quantification have been developed and extensively optimized over the past decades^2–6^, leaving little room for improvement in this area, for completeness, we also assessed the performance of GW distances in relation to state-of-the-art methods.

For that purpose, we considered the problem of identifying RdRp core domains from previously unseen ribovirus phyla. We supplemented our RdRp dataset with 300 computationally predicted structures from non-RdRp sequences (“decoys”), including eukaryotic, bacterial, retroviral, and DNA-viral protein fragments that clustered at 97% identity with true RdRp core domains^48^ (Supplementary Table 2). We then trained a nearest-neighbor classifier on a subset of RdRp core domain structures and decoys, and tested its ability to distinguish between decoys and RdRp core domain structures from individual phyla not included in the training data (Methods). In this evaluation, the embedding space underlying the nearest neighbor classifier was constructed using either GWProt, Foldseek^6^, or TM-align^5^.

GWProt achieved the highest overall performance among the three methods, with a mean MCC of 0.953 compared to 0.927 and 0.922 for TM-align and Foldseek, respectively, although it was not the best-performing method within any individual phylum (Supplementary Fig. 3). This result was primarily driven by the lower performance of Foldseek and TM-align in identifying Lernaviricota and Negarnaviricota riboviria, respectively. We attribute the greater stability of GWProt across different phyla to its solid mathematical foundation, suggesting that combining GWProt with TM-align and Foldseek provides the most robust framework for identifying previously unseen phyla. Indeed, a majority-vote classifier integrating the three methods achieved an average MCC of 0.981, substantially overperforming any individual algorithm (Supplementary Fig. 3).

### Local geometric distortions across the human proteome reveal the structural lexicon underlying functional domains

Protein functional domains are composed of recurrent structural elements at different hierarchical levels. While the classification of functional domains is well established (e.g., via the CATH^12^ and SCOP^13^ databases), there is no comprehensive and systematic classification of the smaller structural motifs that make up functional domains beyond secondary structures, such as α-helices, β-sheets, and combinations of them. We therefore applied GWProt to systematically identify and characterize small structural motifs across protein domains catalogued in the CATH database.

For each CATH homologous superfamily with at least 5 elements (*n* = 552 superfamilies containing 10,139 domains), we computed pairwise GW correspondences and the local geometric distortions between the predicted structures of every element in the superfamily (Methods). We divided each domain into a set of short structural fragments or “clips” that are shared within its superfamily by removing regions with high average local geometric distortion (Fig. 3A). This procedure yielded a total of 66,937 clips. The average length of each clip was 22.5 residues, and each domain contained, on average, 6.6 clips (Figs. 3B and C). Most clips contained α-helices, β-sheets, or both, whereas only 2.3% lacked these elements (Fig. 3D).

**Figure 3.**
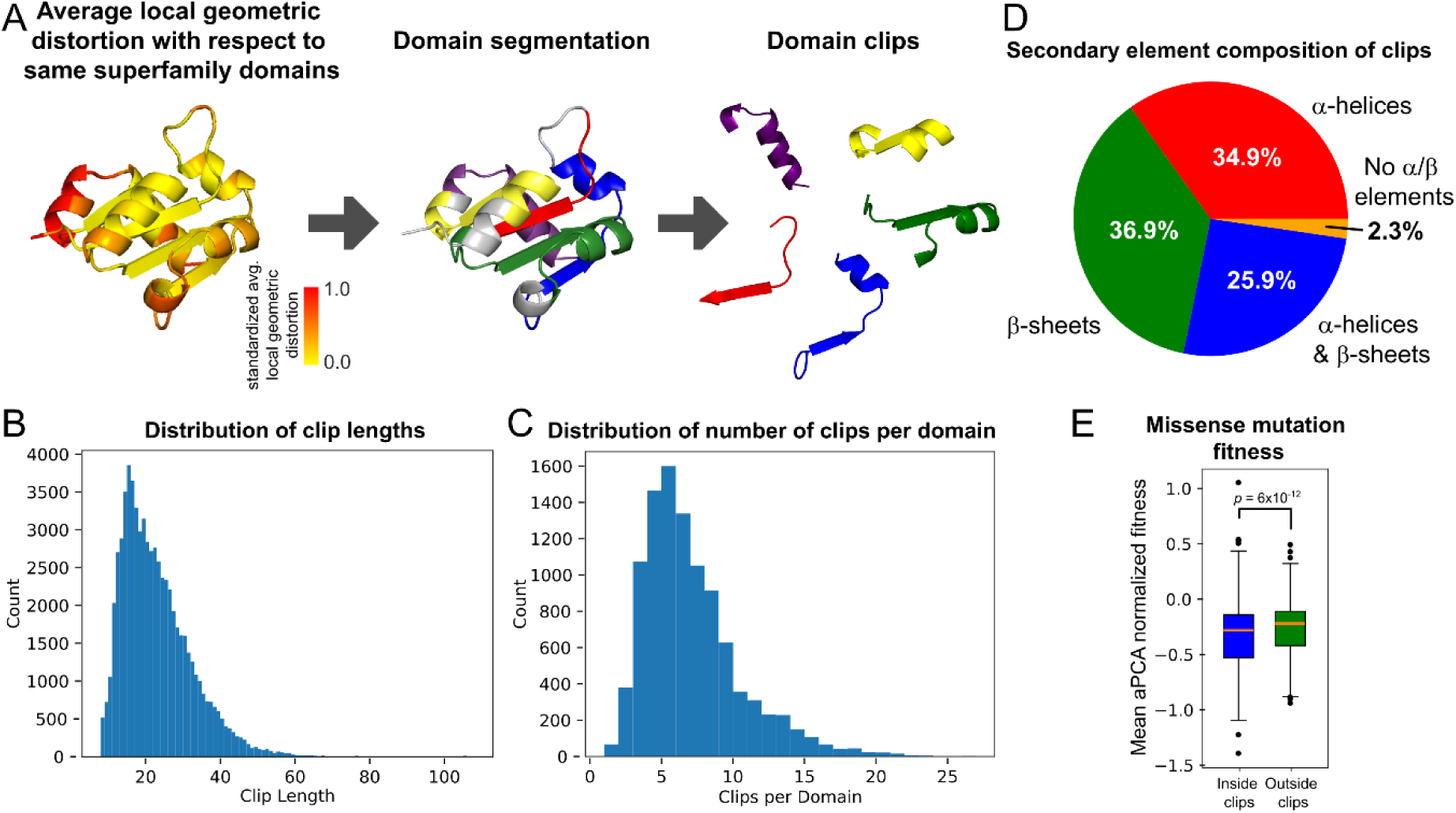
Structurally conserved fragments across human functional protein domains. **A)** Schematic of the approach. Each protein domain in the CATH database is aligned with all other domains belonging to the same homologous superfamily using GWProt, and the average local geometric distortion of each residue in the domain is computed. The average local geometric distortion is then used to segment structurally conserved domain clips. **B)** Distribution of domain clip lengths, measured in number of residues. **C)** Distribution of the number of clips per domain. **D)** Secondary structural element composition of domain clips. **E)** Box plot summarizing the distribution of mean normalized fitness of missense mutations inside domain clips and outside domain clips, but within the domain. aPCA: abundance protein fragment complementation assay.

Clips were significantly depleted of common (MAF ≥ 1%) missense single-nucleotide polymorphisms (SNPs) compared to other regions of the functional domain, suggesting that they are more likely to contain key active or structural sites within the functional domain (odds ratio = 0.93, Fisher’s exact test *p*-value = 0.004). Consistent with this, comparison with a site-saturation mutagenesis screen of 500 protein domains^62^ revealed that missense mutations within the clips are more detrimental to protein stability than those outside the clips but still within the functional domain (Fig. 3E; fold change in mean normalized fitness = 0.8, Wilcoxon rank-sum test *p*-value = 6 x 10^-12^). Thus, local geometric distortion captures information relevant to protein stability and function.

To determine whether structural clips are unique to individual homologous domains or represent a shared structural lexicon across domain families, we computed pairwise GW distances between the 66,937 clips and clustered the resulting space using density-based clustering^63^, yielding 234 clusters of structurally homologous clips, or “structural motifs” (Fig. 4A, Supplementary Fig. 4, and Supplementary Table 3). As expected, the space of clip structures was relatively continuous, and only 35% of the clips were assigned to a cluster by the density-based clustering algorithm. Each cluster contained an average of 101 clips (Fig. 4B), and clip lengths within clusters were nearly unique, varying by at most three amino acids. Notably, each motif was found on average across 4.6 architectures, 18.8 topologies, and 27.7 homologous superfamilies of the CATH domain classification, indicating that these structural motifs are widely shared across the human proteome (Fig. 4B). Moreover, pairs of structural motifs often significantly co-occurred within the same functional domain (Fig. 4C; Fisher’s exact test FDR < 0.05), underscoring the combinatorial nature of the structural lexicon that underlies functional domains.

**Figure 4.**
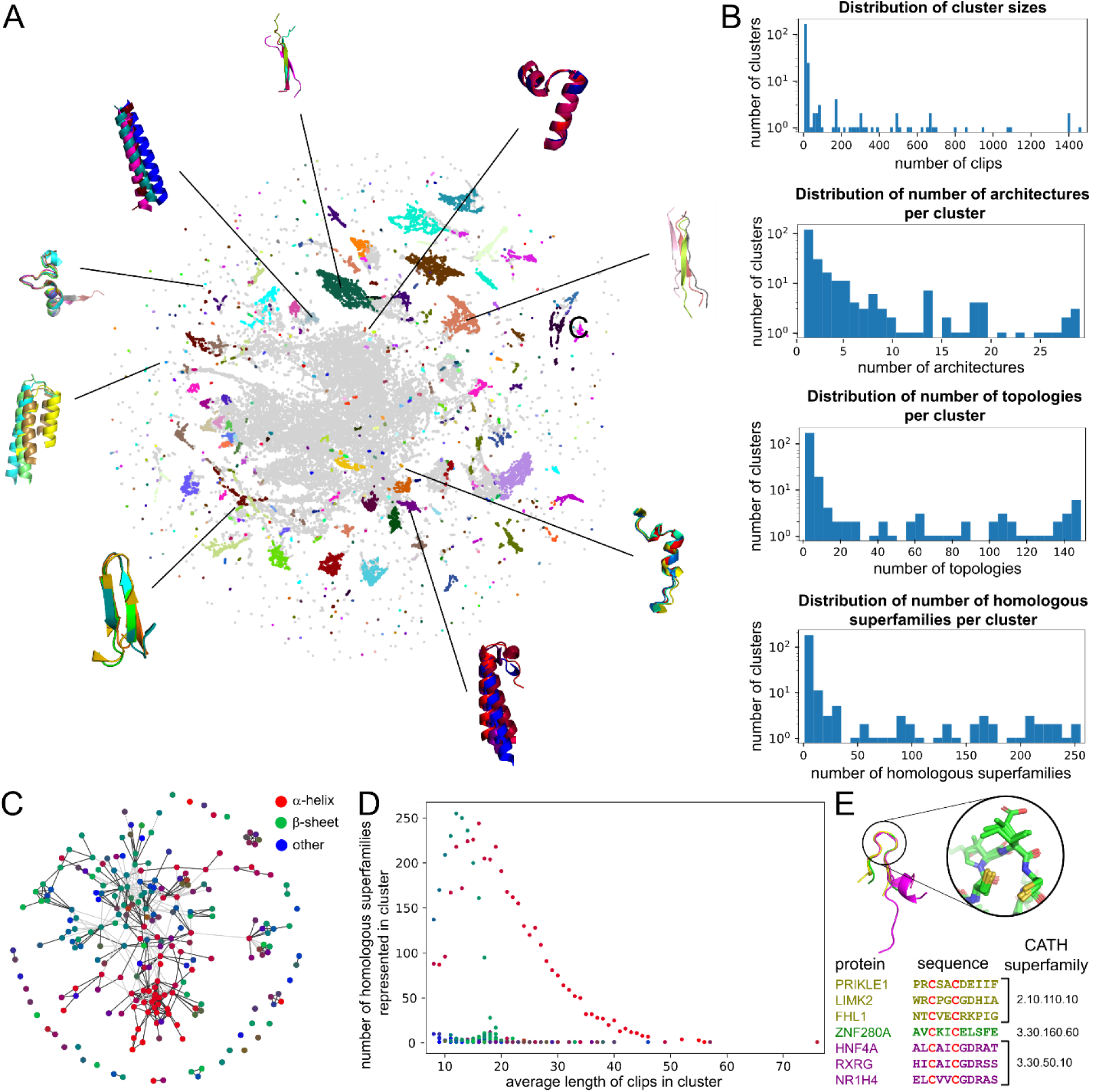
Structural motif analysis across human protein functional domains. **A)** UMAP embedding of structurally conserved protein short fragments across human functional domains. The space was clustered using DBSCAN, and clusters are shown in the UMAP, with representative fragments from selected clusters shown for reference. **B)** Distributions of the number of structural clips per cluster and the number of CATH architectures, topologies, and homologous superfamilies represented in each cluster. **C)** Network showing the co-occurrence of structural motifs within the same functional domain. Each node represents a cluster of structural clips, and edges represent significant co-occurrence (Fisher’s exact test FDR < 0.05) of two motifs within the same functional domain, with edge thickness proportional to the odds ratio (one-sided Fisher’s exact test). Nodes are colored by the proportion of residues in α-helices (red), β-sheets (green), or neither (blue). **D)** Number of CATH homologous superfamilies represented in each structural motif as a function of the average motif length. Points are colored by the secondary structure element composition, as in panel (C). **E)** Examples of short clips belonging to the same structural motif (cluster ID 93), consisting of a small unstructured loop stabilized by a CXXC disulfide motif. The clips span seven proteins and functional domains from three different homologous superfamilies (represented with different colors). Protein names, amino acid sequences, and CATH homologous superfamily identifiers are also shown for reference.

Visualizing the relationship between average clip length, number of homologous superfamilies, and proportion of residues in α-helices and β-sheets for each cluster revealed three main types of structural motifs (Fig. 4D). The first type consists of motifs composed primarily of α-helices, often characterized by bends and diverse terminal loops. These span a wide range of lengths, with shorter motifs being more commonly observed across different CATH homologous superfamilies. The second type includes short motifs (< 20 residues) primarily composed of β-sheets, which are found in a large number (>75) of CATH homologous superfamilies. Finally, the third type consists of short motifs that appear in fewer than 25 homologous superfamilies. This type includes a great variety of β-hairpin structures as well as other structural motifs such as small loops stabilized by CXXC disulfide motifs (two cysteines separated by two other residues) (Fig. 4E).

Taken together, these results reveal a structural lexicon of small motifs shared across distinct functional domains, comprising not only α-helices and β-sheets but also other conserved small structural elements.

### Certain structural motifs show enrichment for pathogenic missense variants independent of domain context

To evaluate the functional relevance of short structural motifs independently of the functional domains in which they occur, we analyzed the positions of 11,130 missense single-nucleotide variants (SNVs) classified as pathogenic or likely pathogenic in the ClinVar database^64^ across the 66,937 clips. These variants were more abundant in clips belonging to the 234 structural motifs (odds ratio = 1.26, Fisher’s exact *p*-value = 10^-24^ after adjusting for clip length).

When examining enrichments in individual motifs, we identified 10 structural motifs that were significantly enriched for pathogenic variants after accounting for clip length and CATH homologous superfamily membership as covariates (permutation test FDR < 0.05; Supplementary Table 4). While some of these enrichments were driven by specific proteins (FOXG1, LDLR, and RPGR) or small groups of closely related proteins (kinases, tubulin β-chains, UDP-glucuronosyltransferases, and cytochrome P450 enzymes), two structural motifs (cluster IDs 11 and 33) showed enrichment of pathogenic SNVs across structurally homologous clips from many unrelated proteins and homologous superfamilies. These motifs consisted of 12-amino-acid β-strand-like domain clips with repeated cysteine/histidine micro-motifs and 35-amino-acid membrane-associated amphipathic α-helices, respectively (Supplementary Fig. 5), highlighting the broad structural and functional role of these motifs across the human proteome.

Collectively, these findings indicate that, consistent with the depletion of common variation in the structural motifs identified by GW correspondences, variants within these motifs are more likely to have pathogenic effects. Furthermore, the widespread occurrence of some of these structural motifs across the proteome provides a framework for relating pathogenic variants among otherwise seemingly unrelated proteins and functional domains.

## Discussion

Metric geometry is the branch of mathematics concerned with the study and comparison of shapes. Here, we investigated the application of metric geometry, and in particular GW correspondences, to the comparative study of protein structures.

Several aspects make GW correspondences particularly attractive in this context. Being mathematically well-grounded, several theoretical results^28–30^ ensure convergence, optimality, stability, and other desirable properties, including the definition of true distances, which facilitate robust implementation of downstream analyses and approximations. For example, the triangle inequality imposes constraints on structural pairwise distances that have not yet been computed, allowing one to determine whether their computation is necessary. This can substantially reduce computational cost, since in most applications only distances to the nearest neighbors need to be computed accurately^34^. Moreover, the modular nature of the GW framework provides flexibility that can be tailored to specific applications. For instance, fused versions of GW correspondences^45^ enable the integration of biochemical information into structural alignments, such as isoelectric points, hydrophobicity, or BLOSUM scores. GW correspondences can also be interpreted locally, allowing the identification of structurally conserved regions. In addition, because they do not rely on sequence information, they remain robust to topologically non-trivial sequence rearrangements.

We have demonstrated these properties across various applications, including the analysis of switch regions in the KRAS oncoprotein, the identification of structurally conserved catalytic sites in RdRps, and the classification of short structural motifs in the human proteome. Our results demonstrate the utility and potential of metric geometry, offering comparable, and in some cases superior, results to existing methods, thereby positioning it as a useful and complementary framework for comparative protein structure analyses. Given the maturity of existing methods, built on decades of algorithmic development and optimization, and the rapid evolution of the emerging discipline of applied metric geometry, with extensions and algorithms for computing GW correspondences advancing quickly, we envision metric geometry as a promising new framework for comparative protein structure analyses whose impact will continue to grow in the years ahead.

Our work has also revealed some of the current limitations in applying metric geometry to protein structure analysis. Specifically, the use of balanced versions of GW correspondences that do not allow for partial matchings poses challenges when comparing proteins that differ substantially in size or contain insertions or deletions of small fragments. While unbalanced GW formulations that address this limitation have been developed^40,46^, their computational cost remains high, limiting their efficient application to thousands of proteins within reasonable timeframes. We anticipate that forthcoming advances in applied metric geometry will help overcome these limitations, broadening its impact on comparative protein structure analysis.

## Methods

### Protein structural alignment with Gromov-Wasserstein correspondences

We define a correspondence between proteins 𝑋 and 𝑌 as a 𝑛 × 𝑚 matrix 𝑇 with entries 𝑇_𝑖,𝑘_ ∈ [0,1] such that ∑_𝑖_ 𝑇_𝑖,𝑘_ = ∑_𝑘_ 𝑇_𝑖,𝑘_ = 1, where 𝑛 and 𝑚 are the number of residues in 𝑋 and 𝑌, respectively. Given a correspondence 𝑇, we define its GW cost as

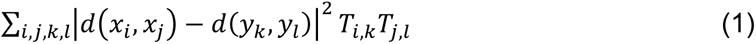

where 𝑑(𝑥_𝑖_, 𝑥_𝑗_) and 𝑑(𝑦_𝑘_, 𝑦_𝑙_) are Euclidean distances between α-carbons 𝑥_𝑖_ and 𝑥_𝑗_ in 𝑋, and 𝑦_𝑘_ and 𝑦_𝑙_ in 𝑌, respectively. The GW distance between 𝑋 and 𝑌 is then given by^29^

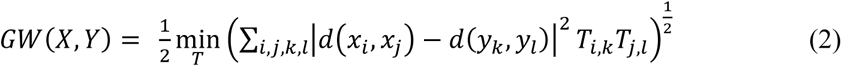

where the minimum is taken over the space of possible correspondences. This quantity satisfies the axioms of a metric space^29^.

Let 𝛿(𝑥_𝑖_, 𝑦_𝑘_) be a non-negative, symmetric function that quantifies the difference between the biochemical properties of any two amino acids. This function can represent a difference in scalar features like isoelectric point, hydrophobicity, or solvent accessible surface area; or pairwise measures, such as the Grantham distance or BLOSUM-based distances. We define the fused GW (FGW) distance as^45^

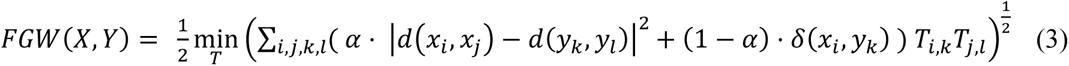

where the parameter 𝛼 ∈ [0,1] controls the relative contribution of the geometric and biochemical distortions. Setting 𝛼 = 1 recovers the standard GW distance, while 𝛼 = 0 corresponds to alignment based solely on biochemical data, ignoring geometric constraints. If 𝛿 defines a metric, then the FGW also defines a metric for a fixed value of 𝛼.

To solve the non-convex optimization problems (2) and (3), we use the network simplex algorithm, as implemented in the Python Optimal Transport (POT) library or our own implementation in the CAJAL package^34^.

Given a correspondence 𝑇 between two protein structures 𝑋 and 𝑌, we define the local geometric distortion of a C_α_ atom in 𝑋 as its contribution to the GW cost function. Specifically,

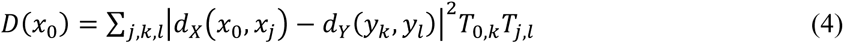

It follows that ∑_𝑖_ 𝐷(𝑥_𝑖_) = ∑_𝑘_ 𝐷(𝑦_𝑘_) = 4 𝐺𝑊(𝑋, 𝑌)^2^. Similarly, in the case of FGW, we define

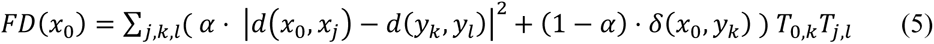

and ∑_𝑖_ 𝐹𝐷(𝑥_𝑖_) = ∑_𝑘_ 𝐹𝐷(𝑦_𝑘_) = 4 𝐹𝐺𝑊(𝑋, 𝑌)^2^. To the best of our knowledge, this pointwise use of GW costs to quantify the geometric distortion of a point 𝑥_0_ when mapped to 𝑌 via a correspondence 𝑇 is novel.

When comparing 𝑁 protein structures in an all-vs-all fashion to identify local geometric distortions, we first perform all 𝑁(𝑁 − 1)/2 pairwise GW (or FGW) computations. For each pair of structures, we store the GW distance and correspondence, and the local geometric distortions.

For a given protein structure, each C_α_ atom accumulates 𝑁 − 1 local geometric distortion values (one from each comparison with the other protein structures). We average these values (in some cases, using a weighted average, as described below) and then transfer the averages across the protein structures using the GW correspondences. Explicitly, the transferred average local geometric distortion for C_α_ atom 𝑥_𝑖_ in 𝑋 from 𝑌 is given by 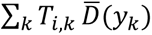, where 𝑇 is the correspondence between 𝑋 and 𝑌, and 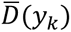 is the averaged local geometric distortion for C_α_ atom 𝑦_𝑘_ in protein 𝑌. This yields 𝑁 − 1 transferred average local distortions per C_α_ atom, which can again be aggregated. This double-averaging procedure substantially enhances the ability to capture local structural variation across the ensemble of protein structures by incorporating composed correspondences.

For visualization, we normalize the local or average local geometric distortion within each protein structure such that the minimum value is 0 and the maximum is 1.

### Identification of KRAS switch regions

We downloaded the PDB files of 54 KRAS protein structures from the RCSB Protein Data Bank^8^, selecting only those determined by X-ray crystallography and containing at most 3 non-synonymous single-nucleotide variants relative to the UniProt^65^ canonical sequence (accession P01116-1) within residues 2-162 (Supplementary Table 1). We restricted our analysis to this sequence region. For PDB files containing multiple KRAS chains, each chain was extracted and saved as a separate structure.

As there is no consensus on the exact boundaries of the switch regions^47^, we adopted residues 30-40 for switch I and 60-72 for switch II as our working definitions.

We aligned all structures using GWProt and computed the average local geometric distortion for each C_α_ atom, as described in subsection “*Protein structural alignment with Gromov-Wasserstein correspondences*” of the Methods section. We computed a single average local geometric distortion for each C_α_ atom, without transferring across structures and second averaging, since the high structural homology made this unnecessary.

To evaluate the predictive power of the averaged local geometric distortion in identifying switch regions, we treated regions with distortion levels exceeding a threshold as predicted switch sites, and calculated precision-recall curves for each protein by varying this threshold.

### Identification of functional sites in RdRps

We considered the computationally predicted structures of the core domain from 2,777 riboviral RdRps with amino acid sequences deposited in GenPept (Supplementary Table 2), out of which 2,739 belonged to one of 14 classes containing at least 10 representatives. Structures were predicted from sequences using AlphaFold^10^ and trimmed to a region of ∼500 residues corresponding to the central core of the RdRp domain^48,49^.

We computed pairwise GW correspondences between the RdRps using GWProt, as described in subsection “*Protein structural alignment with Gromov-Wasserstein correspondences*” of the Methods section. To reduce computation time given the large number of pairwise comparisons (∼3.8 million), we downsampled each structure to 200 evenly spaced residues.

Additionally, we computed pairwise GW correspondences without downsampling for 97 randomly selected RdRps from the same dataset. For these proteins, we identified the locations of the A, B, and C regions using Palmscan^48^ with default parameters, and visually confirmed them in Pymol. We use the GW correspondences to compute local geometric distortion using the double averaging approach described in subsection “*Protein structural alignment with Gromov-Wasserstein correspondences*”. To assess the predictive power of the averaged local geometric distortion for identifying A, B, and C sites, we treated regions with distortion levels below a threshold as predicted A, B, or C sites, and calculated precision-recall curves for each protein by varying this threshold.

Finally, we repeated the analysis on the same set of 97 RdRps using FGW correspondences instead of GW, with parameter 𝛼 = 0.2 and 𝛿 defined as the difference between the standardized consensus hydrophobicity level of the two residues^66^.

### Evaluation of GW distances for the identification of novel RdRps

We considered the computationally predicted folds of the core domains of 150 randomly selected RdRps from each riboviral phylum (Duplonaviricota, Kitrinoviricota, Lenarviricota, Negarnaviricota, and Pisuviricota) with sequence lengths between 200 and 700 residues (Supplementary Table 2). For Lenarviricota, only 124 predicted folds were available, and all were included. We augmented this dataset with 300 non-RdRp decoy structures that clustered at 97% amino acid sequence identity with bona fide RdRp core domains^48^.

We constructed a protein structure embedding space for this combined dataset using FGW, with parameter 𝛼 = 0.2, 𝛿 defined as the difference between the standardized consensus hydrophobicity level of the two residues^66^, and inter-carbon distances scaled using the square root function. Similar embedding spaces were also generated using TM-align^5^, with parameters `-fast -a T`, and Foldseek^6^, with parameters `easy-search –-exhaustive-search 1`.

For each phylum, we then trained a *k* = 3 nearest-neighbor classifier on each of the three embedding spaces using 10-fold cross-validation. In each iteration, 10% of the decoys and 10% of the RdRps from the focal phylum were used as test data, while RdRps from other phyla and the remaining decoys served as the training set. The accuracy of the classifier was quantified for each phylum and embedding space using the MCC.

### Analysis of structural motifs in human protein functional domains

We considered all homologous superfamilies in the CATH database that contained at least 5 human protein domains and obtained their computationally predicted structures from AlphaFold2 as reported by Bordin et al.^67^. For the 78 superfamilies containing over 50 domains, we randomly selected 50 structures. For each domain, we removed residues at the domain termini with predicted Local Distance Difference Test (pLDDT) scores ≤ 75 and excluded domains with fewer than 20 remaining amino acids. After applying these filters, 10,139 computationally folded domains remained.

We computed all pairwise GW correspondences between domains within each homologous superfamily and averaged the local geometric distortions, following the procedure described in the subsection “*Protein structural alignment with Gromov-Wasserstein correspondences*”. To emphasize residues with high local geometric distortion, while keeping distortion values independent of protein size or the number of domains in a superfamily, we averaged the squared distortion values, weighting each domain by the cube of its length and inversely by the total sum of distortion values, and then averaged the transferred distortion values weighted by the length of the domain.

We then computed a rolling average of the local geometric distortion values using a 7-residue window, and selected residues that had an average distortion below the median distortion for the domain (and at most 100,000), except for residues whose average distortion exceeded that of the neighboring 8 residues on each side.

Contiguous residues were grouped into clips. Clips shorter than 8 residues were discarded unless they were within two residues of another clip, in which case the two clips were merged into a single clip. The resulting 66,937 clips were clustered according to their GW distance using the density-based algorithm DBSCAN^63^, as implemented in the Python package scikit-learn, with parameters `leaf_size = 100` and `min_samples = 10`, yielding 23,668 clips grouped into 234 clusters. For each cluster, the proportion of amino acids identified as belonging to α-helices and β-sheets was inferred using the algorithm STRIDE^68^.

For each pair of clusters, we counted the number of domains containing clips from both clusters, from only one of the clusters, or from neither. We then performed a one-sided Fisher’s exact test and corrected for multiple hypothesis testing using the Benjamini-Hochberg procedure. Finally, we constructed an adjacency graph of co-occurrences with q-values < 0.05, weighting the edges by the corresponding odds ratios.

For the analysis of common variation, we annotated common SNPs in the 10,139 CATH domains using SnpEff v5.2f^69^ and the dbSNP database^70^ (build 151), identifying a total of 19,059 residues with common missense SNPs. We then counted the number of residues within clips and outside clips (but still within domains), both with and without common missense SNPs, and performed a Fisher’s exact test to assess the significance of the association between clips and common variants.

To evaluate the stability of structural clips, we calculated the average normalized fitness score for each residue in 341 domains from 203 proteins that overlapped with the site-saturation mutagenesis screen of Beltran et al.^62^. We then assessed the significance of the difference in mean normalized fitness scores between residues in clips and those outside clips (but still within domains) using the Wilcoxon rank-sum test.

### Analysis of pathogenic variants within structural motifs

We retrieved all pathogenic and likely pathogenic missense SNVs from the ClinVar database^64^ (update 2025-08-31) and annotated their position in the clips using SnpEff v5.2f^69^. In total, 11,130 clip positions contained at least one pathogenic or likely pathogenic SNV.

For hypothesis testing, we used the total number of SNV locations within each cluster as our test statistic. We tested clusters with at least 4 SNV locations using a permutation test with 10,000 random permutations. In each permutation, and for each CATH homologous superfamily, we randomly reassigned SNV locations among all clips belonging to domains from that superfamily, with the probability of relocation to a clip proportional to its length. For each cluster, we then calculated the fraction of permutations in which the total number of SNV locations was at least as high as in the unpermuted one. We controlled the false discovery rate using the Benjamini-Hochberg procedure.

### Code availability

The source code and documentation of GWProt are available at https://github.com/CamaraLab/GWProt and https://gwprot.readthedocs.io, respectively.

## Data availability

The computationally predicted structures of the core domain from RdRps and decoys considered in this work are available at https://zenodo.org/records/17468995.

## Acknowledgements

The authors are grateful to Artem Babaian and Robert Edgar for valuable discussions and feedback during the completion of this work, as well as for providing the folds of RdRp core domains and decoys. They also thank Patrick Nicodemus for helpful discussions regarding the implementation of GW computations. P.G.C. acknowledges the organizers of the 21^st^ Annual Workshop at Bellaris, where the idea for this project was conceived. This work was supported by the U.S. National Institutes of Health (NIH) through grant RF1MH130553 from the National Institute of Mental Health (NIMH).

## Author contributions

E. G. and P. G. C. conceived the methodology and wrote the manuscript. E. G. developed and implemented GWProt and performed all the computational analyses. P. G. C. supervised the project.

