## Supplemental Information for "Exploring the Structural Lexicon of the Proteome via Metric Geometry"

### Supplementary Figures

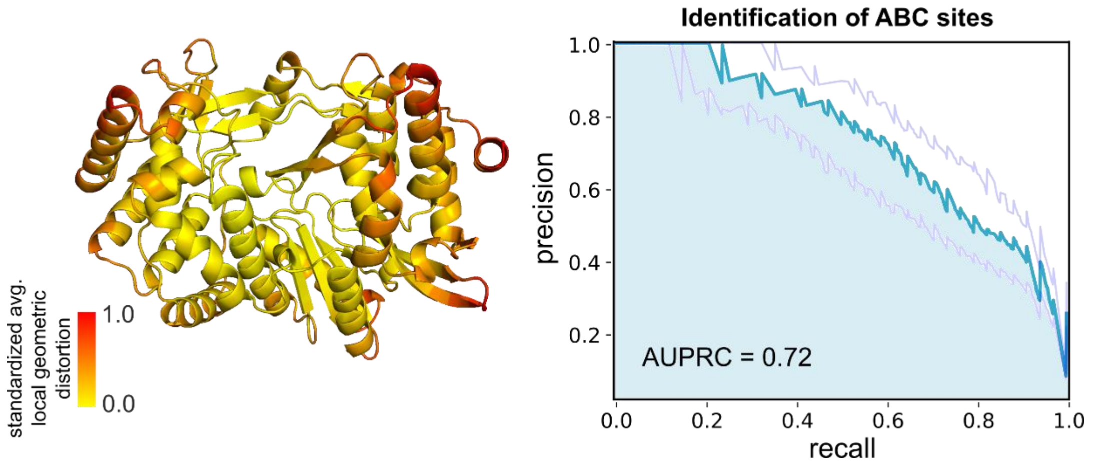


**Supplementary Figure 1. Identification of structurally conserved catalytic sites in RdRps using GW correspondences.** Left: Example of an RdRp core domain from Hepacivirus hominis (GenPept ID AFD18577) colored by average GW local geometric distortion. Regions of low distortion, mostly located at the catalytic core, are structurally conserved. Right: Median, 20%, and 80% percentile precision-recall curves for predicting A, B, or C sites based on average GW local geometric distortion across 97 randomly selected RdRps. AUPRC: area under the precision-recall curve.

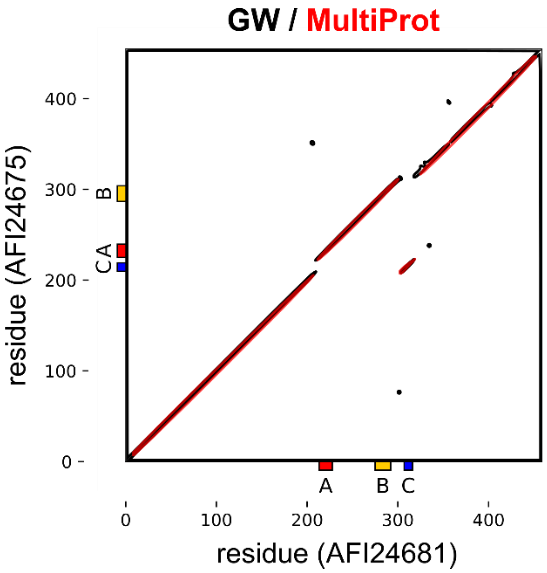


**Supplementary Figure 2. Dot plot visualization of the GW (black) and MultiProt (red) correspondences between two RdRp core domains with ABC and CAB active site orderings.**

| | **GWProt** | **TM-align** | **Foldseek** | Majority-vote |
|---|---|---|---|---|
| Duplornaviricota | 0.963 (0.044) | 0.970 (0.024) | 0.985 (0.022) | 0.990 (0.020) |
| Kitrinoviricota | 0.962 (0.041) | 1.000 (0.000) | 0.995 (0.014) | 0.995 (0.014) |
| Lenarviricota | 0.945 (0.042) | 0.960 (0.053) | 0.680 (0.126) | 0.966 (0.038) |
| Negarnaviricota | 0.936 (0.050) | 0.711 (0.074) | 0.970 (0.033) | 0.971 (0.045) |
| Pisuviricota | 0.957 (0.033) | 0.995 (0.015) | 0.980 (0.024) | 0.985 (0.022) |
| **Average** | 0.953 (0.044) | 0.927 (0.117) | 0.922 (0.136) | 0.981 (0.032) |

**Supplementary Figure 3. Identification of RdRp core domain structures from previously unseen phyla.** Shown are the average 10-fold cross-validation MCC values and standard deviations (in parenthesis) for a $k$ = 3 nearest neighbor classifier trained on structural embedding spaces produced by GWProt, TM-align, and Foldseek. The classifiers were evaluated on the task of distinguishing RdRp core domain structures belonging to phyla absent from the training data from non-RdRp decoys with high amino acid sequence similarity to bona fide RdRp core domains. The MCC of a majority-vote classifier combining the three methods is also shown.

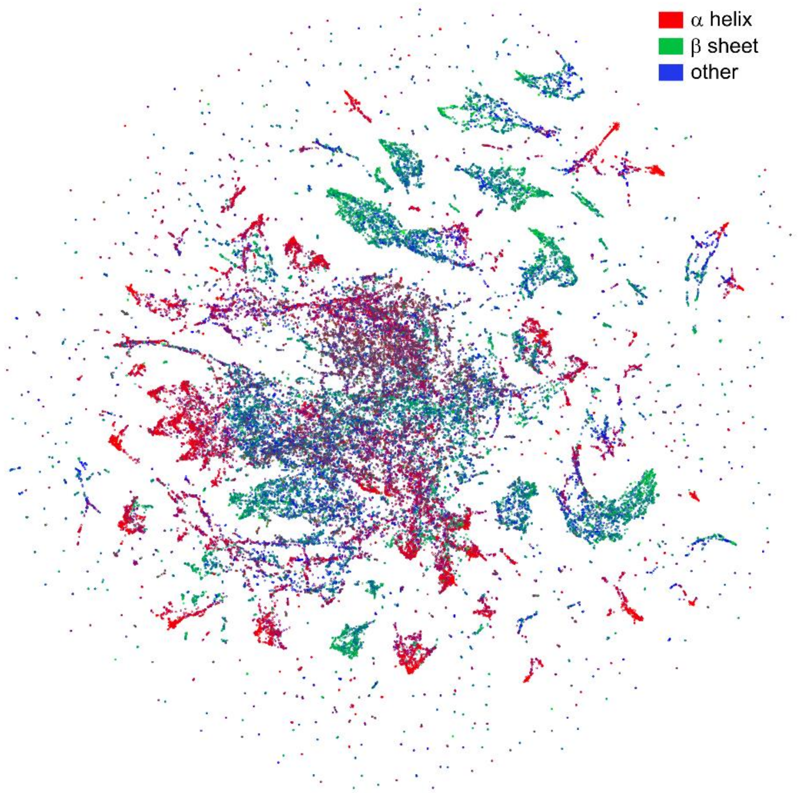


**Supplementary Figure 4. UMAP embedding of the structural space of domain clips, colored according to secondary structure element composition.**

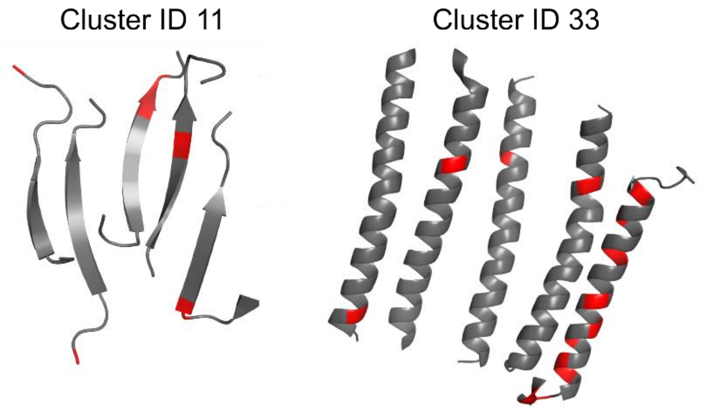


**Supplementary Figure 5. Representative clips from structural motifs 11 and 33.** Positions with known pathogenic or likely pathogenic SNVs are indicated in red.

### Supplementary Tables

**Supplementary Table 1. RCSB Protein Data Bank accession numbers and metadata of the 54 KRAS protein crystallographic structures included in the analysis.** [Provided as a separate file].

**Supplementary Table 2. GenPept accession numbers and taxonomy of the RdRps and non-RdRp decoys included in the analysis.** [Provided as a separate file].

**Supplementary Table 3. List of structurally conserved polypeptides across human functional protein domains.** [Provided as a separate file].

**Supplementary Table 4. Pathogenic missense variant enrichment in structural motifs.** [Provided as a separate file].
